# Triplet Quenching by Active Site Cysteine Residues Improves Photo-stability in Fatty Acid Photodecarboxylase

**DOI:** 10.1101/2025.10.10.681628

**Authors:** Junfeng Ma, Jason M. Kalapothakis, Harry J. Spacey, Linus O. Johannissen, Eugene Shrimpton-Phoenix, Muralidharan Shanmugam, Michiyo Sakuma, Christopher W. Wood, Perdita E. Barran, Derren J. Heyes, Nigel S. Scrutton

## Abstract

Enzyme photobiocatalysis uses light to drive high-energy transformations but is limited by the rarity of photoenzymes. Fatty acid photodecarboxylase (FAP), a recently discovered photoenzyme, enables fatty acid conversion to alkanes/alkenes via excitation of an FAD cofactor, though its poor photostability and photoinactivation has hindered industrial applications. Here, we combine protein engineering approaches with biocatalytic and biophysical techniques, as well as computational chemistry, to demonstrate that additional active site cysteine residues can suppress oxygen-mediated inactivation processes that are driven by the FAD triplet-excited state. We identify a number of positions close to the FAD for cysteine residues that lead to a significant enhancement in activity as a result of an increase in the number of catalytic turnovers and improved photostability. The additional cysteine residues quench the triplet excited state of the FAD cofactor via a proposed proton-coupled electron transfer mechanism, resulting in lower levels of harmful reactive oxygen species. Our study highlights promising routes to mitigate non-productive, photoinactivation pathways in FAP and informs the rational design of new flavin-based photoenzymes.

## INTRODUCTION

Photobiocatalysis, reactions catalyzed by a light dependent catalyst or photoenzyme, has advanced rapidly in recent years. However, only three naturally occurring classes of photoenzymes have been discovered to date, namely DNA photolyase^1^, protochlorophyllide oxidoreductase^2^ and the recently discovered fatty acid photodecarboxylase (FAP)^3^. Of these three, only FAP has demonstrated the potential for synthetic applications^4^. Inspired by these natural photoenzymes, thermally active flavin-binding enzymes have been repurposed and redesigned, and protein scaffolds have been engineered to bind synthetic chromophores, to catalyze new-to-nature light-driven chemistry^5-9^. Nevertheless, photoinactivation and photostability presents a major barrier for their more widespread use in industrial applications and has limited the rational or *de novo* design of more efficient photoenzymes.

The FAP enzyme from the microalga *Chlorella variabilis* (*Cv*FAP) has proven to be a promising target for sustainable biofuel production^10^ since it uses abundant fatty acids as a substrate. Additionally, biotransformations can be extended by incorporating a further step, enabling the use of triglycerides, which are accessible from renewable feedstocks^11-12^. The reaction scope of FAP has been expanded by using modified versions of the enzyme to catalyze the separation of racemic mixtures and the synthesis of other various compounds, such as long-chain secondary alcohols, esters, and amines^4, 13-16^. More recently, building on the pioneering flavoenzyme engineering studies of Hyster^6, 17-21^ and Zhao^7, 22- 23^, Yang and co-workers have demonstrated that FAPs can be adapted to drive new stereoselective radical transformations^24^.

Biophysical and structural studies have provided a detailed understanding of the mechanism of photocatalysis in FAP^3, 25-26^. Fatty acid (FA) substrate binds to the FAP–FAD holoenzyme complex in the oxidized FAD state. Excitation of this enzyme-substrate complex generates a FAD singlet excited-state ^1^FAD^*^, which abstracts an electron from FA to form a flavin semiquinone FAD^•−^ and a high energy FA^•^ radical. The FA^•^ radical spontaneously decarboxylates to release carbon dioxide and the resulting alkyl radical is converted to the alkane product via electron and proton transfer reactions, which are coupled with the reoxidation of the FAD semiquinone. These latter steps proceed through a novel, red-shifted FAD intermediate and are proposed to involve an active site cysteine, Cys432, and arginine residue, Arg451^25-26^. By developing a continuous spectroscopic assay for the detection of the CO_2_ product it was recently shown that violet light is more efficient than blue light at driving this reaction chemistry, due to the formation of higher populations of the FAD excited state^27^.

Despite significant advances in our mechanistic understanding of the catalytic cycle of FAP, photostability remains a major problem^25, 28^ and the exact mechanism of photoinactivation is still unclear. Photoinactivation is thought to be driven by the triplet excited state of the FAD, which is formed as an off-pathway process upon illumination in the absence of substrate (Figure 1A). The FAD triplet reacts with molecular oxygen and leads to the formation of reactive oxygen species (ROS)^29^. The ROS are able to react with active site residues, leading to formation of free radicals and irreversible damage to the protein, most likely via cross-linking, adduct formation and backbone cleavage that cause misfolding, aggregation, degradation or cofactor release^27, 30-31^. Attempts to circumvent photoinactivation have included using a significant excess of FA substrate^32^ or decoy molecules^33^ to keep the enzyme catalytically ‘busy’, performing reactions under anaerobic conditions^29^ or using pulses of low-intensity violet light^27^ to provide an efficient route for productive catalysis whilst minimizing inactivation. However, none of these tackle photostability directly and are unlikely to be long-term solutions to the photoinactivation problem. In non-enzymatic systems, L-cysteine has previously been reported to efficiently quench the triplet excited state of riboflavin in solution^34^ and may provide a plausible route to minimize the harmful off-pathway inactivation in FAP.

**Figure 1.**
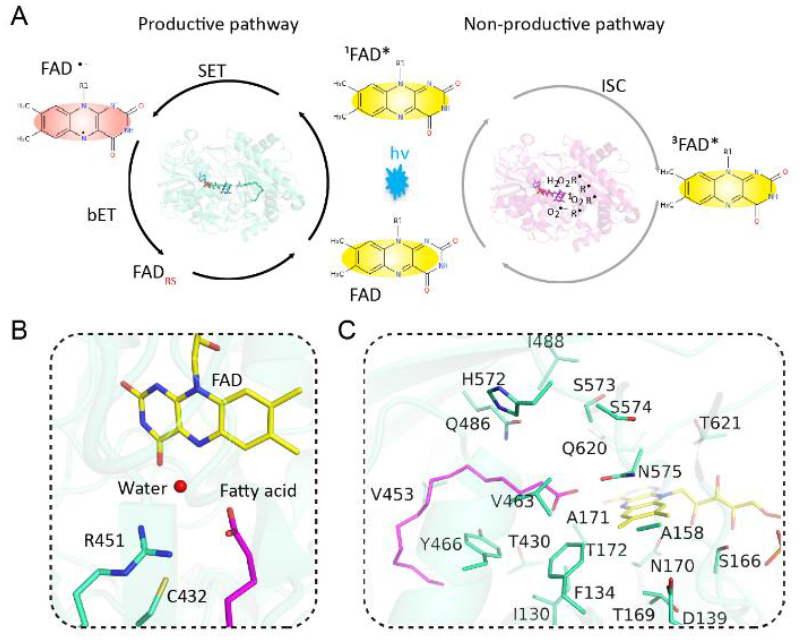
Structural and mechanistic rationale of photocatalysis and photoinactivation in FAP. (A) Productive photocycle (left panel) proceeds via an electron transfer (SET) from the fatty acid substrate to the singlet excited state of the FAD cofactor, followed by back electron transfer (bET) to yield reoxidized FAD via a red-shifted intermediate (FAD_RS_). There is also a non-productive pathway in the absence of substrates, which proceeds via the FAD triplet state through intersystem crossing (ISC) from the singlet state (right panel), leading to photoinactivation of the enzyme. (B) The active site of FAP showing the position of the catalytic residues, Cys432, and Arg451, relative to the FAD cofactor and fatty acid substrate. (C) The position of the residues selected in the current work for protein engineering studies (shown as lime green sticks). The fatty acid substrate is shown in magenta and FAD cofactor in yellow. PDB: 5NCC.

Using protein engineering approaches we have identified a number of additional active site Cys residues, located close to the isoalloxazine moiety of the FAD cofactor, that cause a significant enhancement in product yield. Detailed kinetic and biophysical analyses show that the additional Cys residues quench the triplet excited state of the FAD, leading to a major reduction in photoinactivation. QM/MM studies reveal triplet quenching proceeds via a proton-coupled electron transfer (PCET) between the FAD and Cys residues, which allows the enzyme to catalyze a significantly higher number of turnovers prior to complete inactivation. Taken together, our work provides new insights into the mechanism of photoinactivation in FAP and highlights a promising approach to improve photostability in FAP and other flavoenzyme systems to further their suitability for industrial applications.

## RESULTS AND DISCUSSION

### Role of Cys and Arg residues in the active site of FAP

In FAP, Cys432 and Arg451 have been shown to play pivotal roles in the FAP photocycle (Figure 1B), with both active site residues strictly conserved among FAP-like proteins^35^. The FAP variants C432S, R451A and R451K exhibit little or no catalytic activity compared to wild-type FAP^25^. In order to explore whether the positioning of these essential residues can influence photocatalysis in FAP we have introduced Cys or Arg residues at different positions around the active site in both C432S or R451A variants (Figure 1C and Table S1). We found only a single variant, C432S/A171C, that could recover any decarboxylation activity, but this variant still remains a lot less active than wild-type FAP (3% activity). Hence, the exact positioning of these catalytic arginine and cysteine residues in the active site is crucial for enzyme activity, highlighting their importance in the catalytic mechanism of the enzyme.

The essential nature of Cys432 and Arg451 in the photochemical mechanism of FAP prompts the question whether variants with additional Cys or Arg residues can improve the catalytic activity of FAP? To this end, we have generated a number of variants that contain double Cys or Arg residues in the FAP active site. Although none of the variants with an additional Arg residue gave rise to any efficient photoenzymes (Table S2), there were several double-Cys variants that produced higher yields of pentadecane product than wild-type (Figure 2A). Three variants with the highest product yields, A158C, A171C and G622C, all contain additional Cys residues that are located near the isoalloxazine ring of the flavin cofactor (Figure 2B). The enhanced activity was maintained after purification of these enzymes (Figure 2C). Hence, they were selected for more detailed activity and biophysical studies to elucidate the mechanistic origin of the enhanced product yields.

**Figure 2.**
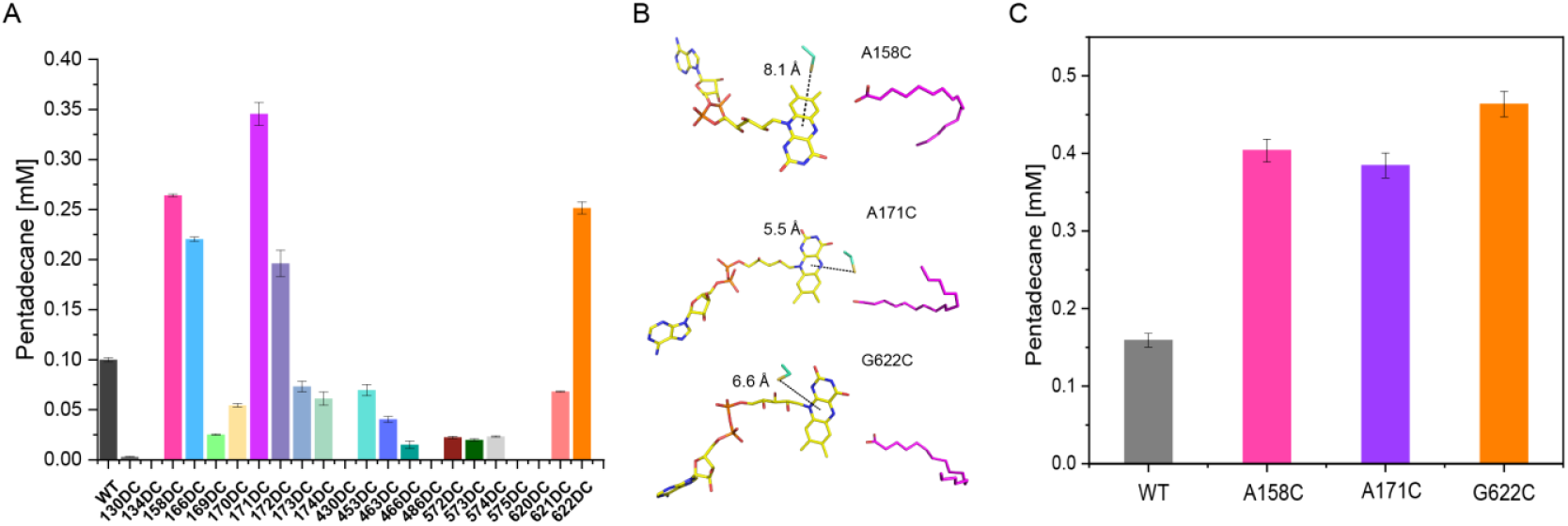
Initial screen of catalytic activity of double cysteine FAP variants. (A) Decarboxylation activity of double cysteine active site variants of *Cv*FAP using cell crude lysate based on standard curve in Figure S2. 1 μM crude enzyme (determined from SDS-PAGE analysis – see Methods section) and 500 µM palmitic acid was illuminated overnight at room temperature at 350 rpm with 1000 μm m^- 2^s^-1^ blue light. Three of the variants displayed highest enhanced activity compared to WT. Other variants showed no activity as shown in Table S2. (B) Architecture of active site of the top three performing variants, showing the distance between sulfur of each cysteine to the center of the isoalloxazine ring. (C) Activity of the top three variants using 1 μM purified enzyme under identical conditions to panel A.

### Kinetics of competing photoinactivation and productive photochemistry in FAP

In order to explore whether the increased product yields in the selected variants were caused by higher rates of catalytic activity or improved photostability of the enzyme we measured the rates of photocatalysis using our recently developed real-time enzyme-coupled assay to directly detect the production of CO_2_^27^. The catalytic rates of A158C and A171C are much lower than that of the wild type (0.52 s^-1^), whereas G622C shows a rate comparable to the wild type, suggesting that any improvements in product yield are not caused by an enhancement in catalysis (Figure 3A).

**Figure 3.**
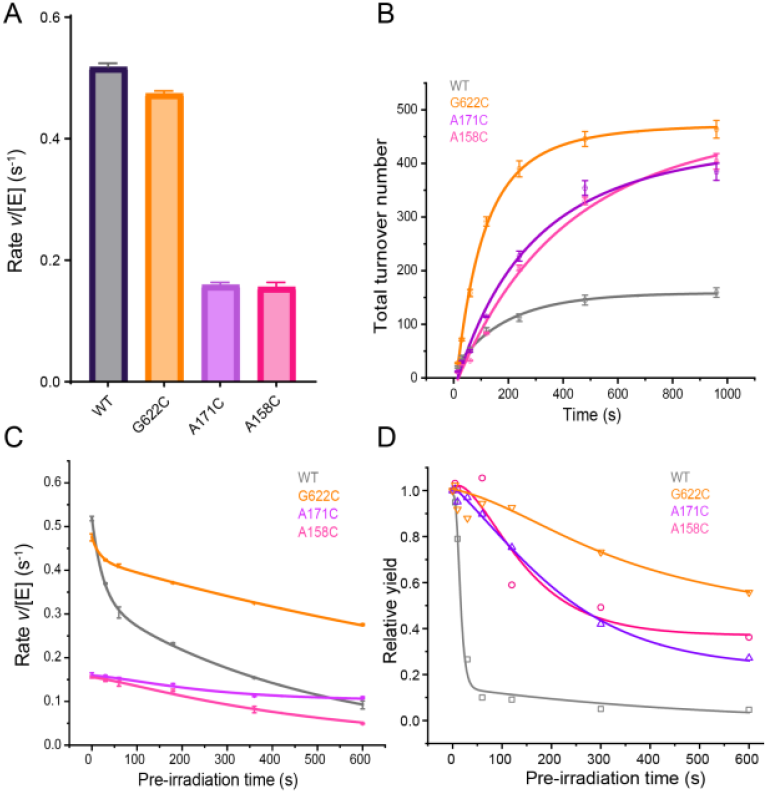
The activity and photostability of wild-type and double Cys variants of FAP. (A) The activity of wild-type and double Cys variants of *Cv*FAP was measured using a real-time stopped-flow coupled assay at room temperature. 5 μM purified enzyme and 0.2 mM palmitic acid substrate were used, and initial apparent reaction rates were determined by measuring the gradient at the early linear region of the traces. The background rate from the assay (when performed in the dark) was subtracted from each measurement to give a final rate. (B) Kinetic trace of pentadecane production vs. reaction time was measured using a gas chromatography assay using 500 μM palmitic acid substrate and 1 μM purified enzyme at room temperature, 350rpm, and 455 nm of 1000 μm m^-2^s^-1^. Experiments were repeated in triplicate and the error bar represents the standard deviation of the measurements. Data was fitted to Eq. 4. (C) Use of the enzyme-coupled assay to measure *Cv*FAP residual activity after various times of pre-illumination with 100 μmol m^-2^s^-1^ blue light on substrate-free enzymes in aerobic condition and room temperature. 5 μM purified enzyme was used, 0.2 mM palmitic acid substrate was added to the reaction mixture after pre-irradiation. Initial apparent reaction rates were determined by measuring the gradient at the early linear region of the traces. The background rate from the assay (when performed in the dark) was subtracted from each measurement to give a final rate. Data points were fitted with Eq. 2d and 3d. (D) Pentadecane yields, detected by gas chromatography with flame ionization, after approximately 30 mins of reaction, after pre-irradiation of substrate-free enzymes for time intervals shown, with a light intensity of 100 μmol m^-2^s^-1^ blue light. 1 μM purified enzyme was used, 0.5 mM palmitic acid substrate was added to the reaction mixture after pre-irradiation and the reaction was performed overnight (∼18hrs) under 350rpm, room temperature and blue light of 1000 μmol m^-2^s^-1^. Data is interpreted with Eq. 5 using rate coefficients from the data shown in 3B.

A time-course kinetic analysis of pentadecane formation confirms that the double Cys variants give higher product yields over longer times (Figure 3B) presumably due to improved photostability allowing a much higher number of catalytic turnovers prior to photoinactivation. Given that these traces include both the productive catalytic and non-productive inactivation pathways, modeling of the kinetic processes was employed to explain the observed differences in activity. The model (see methods section) describes how competing photocatalytic and photoinactivation pathways may vary between the variants during the time course of the measurement, taking into account the consumption of substrate over time. In order to isolate and determine the profile of the inactivation pathway, we measured the decay in the initial apparent reaction rates using our stopped-flow enzyme-coupled assay^27^. The wild-type and variant enzymes were preirradiated for various lengths of time, specifically with 0, 30, 60, 180, 360 and 600s of irradiation, and initial rates of reaction were measured using these irradiated FAP enzymes (Figure 3C). The initial rate is given by Equation 1a 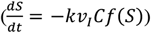, where *C*, the concentration of active enzyme is now a function of the inactivation time. Notably, a single first-order process does not suffice to interpret the observed dependence of reaction rates on pre-irradiation times. Instead, for the wild-type and G622C FAP enzymes there are two measurable decay processes, a fast and a slow photo- inactivation process. However, the relative proportion of the fast inactivation process is markedly reduced in the G622C variant (10%) compared to the wild-type enzyme (36%).

There are two potential interpretations for the presence of two such inactivation processes in FAP. Firstly, it is possible that there is an oxygen-dependent and oxygen-independent inactivation pathway. However, we observed only minimal photoinactivation under anaerobic pre-irradiation (Figure S3B), suggesting that this is highly unlikely. Alternatively, the data could be explained by the presence of two distinct and slowly interconverting conformations of the enzyme, which have different inactivation rates. Interestingly, in our related work we have indeed observed two distinct conformational ensembles of FAP by using mass spectrometry^36^. In contrast to the wild-type and G622C FAP enzymes, there is only a single slow photo-inactivation process in the A158C and A171C variants and the fast kinetic phase is absent.

The differences in photoinactivation rates can be used to interpret the kinetic traces of pentadecane production over time (Figure 3B and Table 1) and relevant parameters can be extracted (Table 1). The suppression of the rapid photo-inactivation process in the double Cys variants appears to be responsible for the observed catalytic efficiency and results in higher product yields. The presence of the initial photo- protective process was further demonstrated by measuring the reaction yields after varying times of pre-irradiation (Figure 3**D**). The product yield after 30 minutes of photocatalysis confirms that the activity of the wild-type FAP decreases rapidly with pre-irradiation time. In contrast, the double Cys variants exhibit a much higher level of photostability, which leads to a significant increase in reaction yields at longer pre-irradiation times. Taken together, this data clearly demonstrates that the double Cys variants are able to undergo many more catalytic turnovers than the wild-type FAP enzyme (2-3 fold higher) as a result of this improved photostability. It is likely that this photoprotective effect will be enhanced at lower light intensities and under different illumination conditions^27^.

**Table 1.** Catalytic turnover and photo-inactivation rate coefficients parameters for *Cv*FAP.

| Variant | Turnover number | Proportion of fast inactivation | Fast inactivation rate coefficient. $k_i[v_i] \text{ s}^{-1}$ | Slow inactivation rate coefficient. $k_i[v_i] \text{ s}^{-1}$ |
| --- | --- | --- | --- | --- |
| WT | 176.9 | 36% | 0.042 | 0.0021 |
| A158C | 413.2 | — | — | 0.0018 |
| A171C | 390.8 | — | — | 0.00073 |
| G622C | 487.4 | 10% | 0.058 | 0.00073 |

### Mechanistic insights into the increased photostability of Cys variants

A rationale for the enhancement in photostability of the Cys variants was investigated. Fluorescence spectra show a decrease in the emission intensity of the three variants compared to wild type FAP, whilst the A158C and G622C variants also exhibited a 10-20 nm blue-shift in the emission maximum of the FAD cofactor (Figure 4A). This suggests that the singlet excited state is quenched in the variants, possibly caused by excited state electron transfer between the FAD cofactor and the Cys residues. It is known that a novel red shifted flavin intermediate (FAD_RS_) is observed during the formation of the final alkane product in the FAP catalytic cycle with a time constant of ∼100 ns^26^. We measured whether the same FAD_RS_ species is formed in the Cys variants using time-resolved laser photoexcitation measurements. A similar increase in absorbance at 515 nm, indicative of the formation of FAD_RS_, occurred with comparable time constants for all of the selected variants. However, the FAD_RS_ species was formed at much lower levels in the A158C and A171C enzymes in accordance with the lower levels of catalytic activity observed in these enzymes (Figure S4).

**Figure 4.**
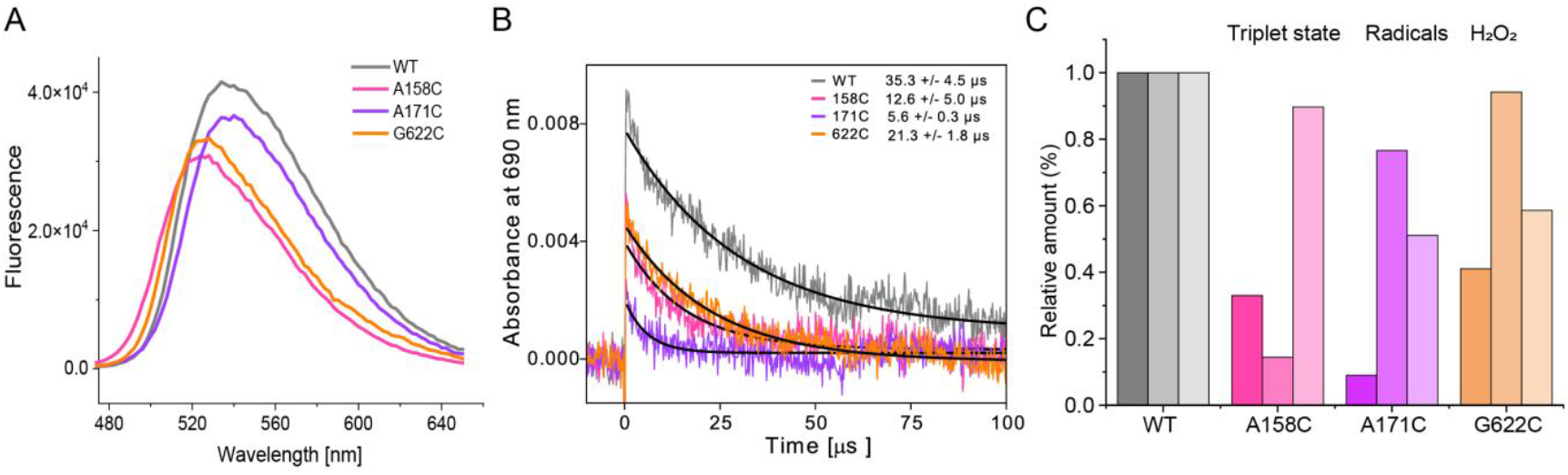
Excited state properties of the FAD cofactor in wild type and double Cys variants of FAP. (A) Steady-state fluorescence of wild type and double Cys variants of FAP after excitation at 450 nm. (B) Kinetic transients showing the decay of the FAD triplet excited state at 690 nm for wild-type and double Cys variants of FAP upon excitation with a 450 nm laser pulse. The time constant of the decay is displayed in the panel. (C) Relative amounts of triplet state, radicals and hydrogen peroxide formed upon illumination with blue light for wild-type and the double Cys variants of FAP. Relative amount of triplet state is obtained from the amplitude in Figure 4B. Relative amount of radicals were quantified according to the integrated area from the EPR signal upon blue light illumination. Relative amount of hydrogen peroxide produced in wild-type and double Cys variants of FAP upon illumination with blue light was measured using a horseradish peroxidase assay. All data is expressed as a relative amount in contrast to wild-type FAP.

Previous studies have shown that FAP readily forms a triplet excited state ^3^FAD^*^ species^3, 25^ in the absence of substrate that is proposed to cause irreversible damage to the protein, likely via reactive oxygen species (ROS)^29^ and radical formation^28^, thereby leading to photoinactivation (Figure 1A). Hence, laser photoexcitation measurements have been used to probe differences in the triplet excited state of the FAD cofactor in each of the Cys variants. Kinetic transients at 690 nm, indicative of the decay of the ^3^FAD^*^ triplet species, were measured under anaerobic conditions to prevent rapid quenching by molecular oxygen. For all three variants the amplitude of the triplet was significantly lower than the wild type enzyme and the triplet state decayed with a shorter lifetime (Figure 4B). These data were used to estimate the relative amount of triplet state over 100 µs for each enzyme, yielding levels of ∼40 %, ∼30 % and ∼10 % respectively for G622C, A158C and A171C (Figure 4C). Therefore, our data suggest that the additional cysteine adjacent to the FAD cofactor inhibits the formation of the triplet state and quenches the triplet state under anaerobic conditions.

The route of photoinactivation in FAP is likely to be via electron transfer between the FAD triplet state and molecular oxygen to produce ROS, which leads to off-pathway radical formation and damage to active site residues^28-29^. We used electron paramagnetic resonance (EPR) spectroscopy to probe radical formation during the photoinactivation pathway for wild-type and variant FAP enzymes (Figure 4C and Figure S5). EPR spectra of samples illuminated with blue light in the absence of substrate showed lower levels of radical species in the Cys variants compared to the wild-type enzyme (Figure 4C). Moreover, we also measured and quantified the H_2_O_2_ produced in the selected FAP enzymes upon illumination with blue light as a proxy for the formation of ROS. Again, the amount of H_2_O_2_ produced was lower in all Cys variants compared to the wild type enzyme (Figure 4C), suggesting that these variants are less susceptible to ROS damage. Taken together, our data provide compelling evidence that the presence of these additional Cys residues in close proximity to the FAD cofactor quenches the triplet state of the FAD, leading to lower levels of ROS that can react with other active site residues to form radicals and irreversible damage to the enzyme.

### PCET between Cys and FAD confers photostability in FAP variants

It has been shown that the triplet state of the flavin cofactor can be quenched by proton-coupled electron transfer (PCET) from nearby tyrosine or tryptophan residues^37^. As cysteine residues can also act as PCET donors^38-39^, we hypothesized that triplet quenching could be facilitated by PCET from C158, C171 or C622 in the double Cys variants, provided that a nearby nucleophile exists that can act as a proton acceptor. MD simulations suggest that the thiol of A158C is hydrogen bonded to the nearby D139 (Figure 5A and 5D), which could potentially act as a proton acceptor, and our QM/MM calculations suggested a barrier of ∼19 kJ mol^-1^ for this reaction. For A171C and G622C, potential proton acceptors are less obvious, although MD simulations suggest that the thiols do transiently form hydrogen bonds with the N5 (A171C) and N1 (G622C) of the isoalloxazine ring of the FAD (Figure 5B and 5E, 5C and 5F). For A171C PCET with proton transfer to N5 was therefore modelled, both *via* a bridging water molecule and with direct H-transfer to N5, since a bridging water molecule is present (>99% of structures) that could also potentially act as a proton shuttle between the thiol and FAD N_5_ (Figure 5E). However, QM/MM calculations suggested that direct PCET is more likely, with a potential energy barrier of ∼53 kJ mol^-1^ compared to ∼74 kJ mol^-1^ for bridged transfer (Figure 5H). For G622C our QM/MM calculations suggest that PCET can occur to the FAD N1 with a potential energy barrier of ∼49 kJ mol^-1^ (Figure 5I). Qualitatively, these barriers are consistent with the observed timescales of triplet quenching (5-20 μs = 60-63 kJ mol^-1^ from the Arrhenius equation). Crucially, the modelled PCET reactions are all exothermic despite different proton acceptors suggesting that this could be a general mechanism for the quenching of excited state triplet flavins.

**Figure 5.**
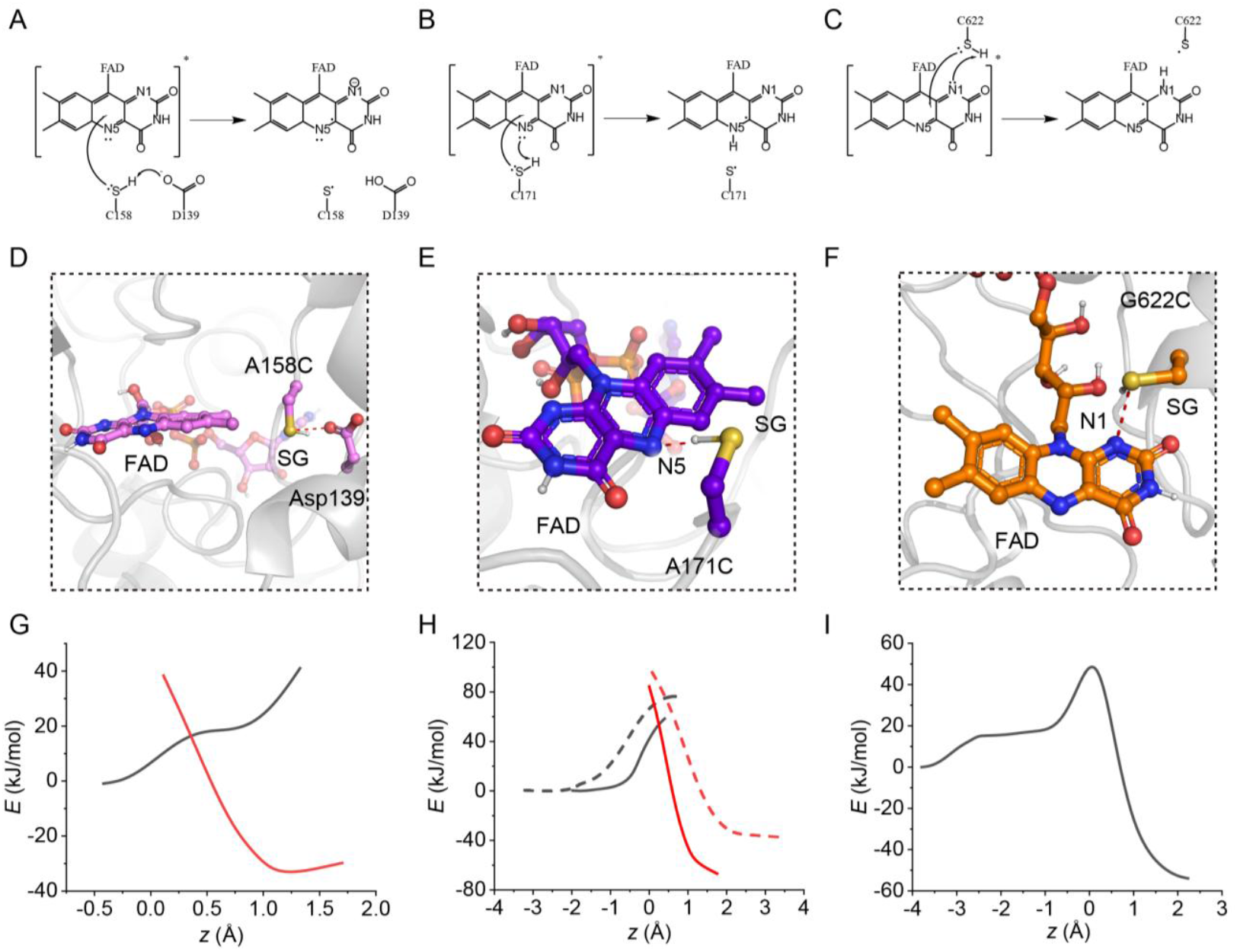
Potential proton acceptors in PCET for *Cv*FAP variants A158C (**A** and **D**) and A171C (**B** and **E**) and G622C (**C** and **F**). Geometries have been taken from MD trajectories. Potential energy plots for PCET from QM/MM calculations for A158C (**G**) and A171C and G622C (**I**). The structures show the reactant and product states for the PCET, and the plots show the potential energy for the electronic states before and after electron transfer as a function of the proton coordinate *z*, defined as the difference in length between the breaking bonds and forming bonds. For A171C this is defined as *z =* [*R*(S_*C171*_-H_*C171*_*)+R(O*_*w*_*-*H_*w*_)] – [*R*(O_*w*_-H_*C171*_)*-R*(N5-H_*w*_)] where S_*C171*_ and H_*C171*_ are the thiol S and H of C171 and *O*_*w*_ and H_*w*_ are the O and transferring H atom of the bridging water molecule; for A158C the proton coordinate is defined as *z =* [*R*(S_*C158*_-H_*C158*_) – *R*(O_*D138*_-H_*C158*_).

## CONCLUSIONS

The industrial application of FAP and other flavin-based photoenzymes has long been limited by their poor photostability under catalytic conditions. In this work, we have addressed this bottleneck through protein engineering, identifying improved FAP variants that incorporate an additional cysteine residue near the FAD isoalloxazine ring, based on the mechanistic understanding of the *Cv*FAP photocycle and flavin dynamics in its excited state. This strategically positioned residue quenches the flavin triplet state via a protoncoupled electron transfer (PCET) mechanism, effectively suppressing oxygen-mediated off-pathway photoinactivation. As a result, these engineered enzymes exhibit significantly enhanced photostability, supporting substantially more catalytic turnovers before complete deactivation. This approach not only advances the practical utility of FAP for light-driven biotransformations but also provides a generalizable strategy for stabilizing other flavin-dependent photoenzymes. The mechanistic insights gained here lay the foundation for the rational design of next-generation photobiocatalysts with improved performance and durability.

## ASSOCIATED CONTENT

### Supporting Information

Details of methods, raw data and control data are shown in figures S1-S7. This material is available free of charge via the Internet at http://pubs.acs.org.

## AUTHOR INFORMATION

### Corresponding Author

- Derren Heyes
- Nigel Scrutton

### Author Contributions

JM, HJS, MiS, MuS and DJH performed experiments and analyzed data. JMK carried out kinetic modelling of the data. LOJ and EPS performed computational chemistry calculations. DJH, CWW, PEB and NSS initiated and coordinated the research program. JM, JMK and DJH wrote the manuscript through contributions of all authors. All authors have given approval to the final version of the manuscript.

### Funding Sources

This work was supported by a Strategic Longer and Larger grant (BB/X003027/1) funded by the Biotechnology and Biological Sciences Research Council (BBSRC) as part of UK Research and Innovation.

## ACKNOWLEDGMENT

The authors thank Dr Kai Wu at Shanghai University of Medicine & Health Sciences for assistance with ROS detection.

## ABBREVIATIONS

FAP: fatty acid photodecarboxylase
FA: fatty acid
ROS: reactive oxygen species
PCET: proton-coupled electron transfer
EPR: electron paramagnetic resonance.

## Notes

### Competing Interest Statement

The authors have declared no competing interest.

## REFERENCES

1. Sancar, A., Structure and function of DNA photolyase. Biochemistry 1994, 33 (1), 2–9.

2. Heyes, D. J.; Zhang, S.; Taylor, A.; Johannissen, L. O.; Hardman, S. J. O.; Hay, S.; Scrutton, N. S., Photocatalysis as the ‘master switch’ of photomorphogenesis in early plant development. Nat Plants 2021, 7 (3), 268–276.

3. Sorigue, D.; Legeret, B.; Cuine, S.; Blangy, S.; Moulin, S.; Billon, E.; Richaud, P.; Brugiere, S.; Coute, Y.; Nurizzo, D.; Muller, P.; Brettel, K.; Pignol, D.; Arnoux, P.; Li-Beisson, Y.; Peltier, G.; Beisson, F., An algal photoenzyme converts fatty acids to hydrocarbons. Science 2017, 357 (6354), 903–907.

4. Zhang, W.; Lee, J. H.; Younes, S. H. H.; Tonin, F.; Hagedoorn, P. L.; Pichler, H.; Baeg, Y.; Park, J. B.; Kourist, R.; Hollmann, F., Photobiocatalytic synthesis of chiral secondary fatty alcohols from renewable unsaturated fatty acids. Nat Commun 2020, 11 (1), 2258.

5. Trimble, J. S.; Crawshaw, R.; Hardy, F. J.; Levy, C. W.; Brown, M. J. B.; Fuerst, D. E.; Heyes, D. J.; Obexer, R.; Green, A. P., A designed photoenzyme for enantioselective [2+2] cycloadditions. Nature 2022, 611 (7937), 709–714.

6. Emmanuel, M. A.; Greenberg, N. R.; Oblinsky, D. G.; Hyster, T. K., Accessing non-natural reactivity by irradiating nicotinamide-dependent enzymes with light. Nature 2016, 540 (7633), 414–417.

7. Huang, X.; Wang, B.; Wang, Y.; Jiang, G.; Feng, J.; Zhao, H., Photoenzymatic enantioselective intermolecular radical hydroalkylation. Nature 2020, 584 (7819), 69–74.

8. Chen, B.; Li, R.; Feng, J.; Zhao, B.; Zhang, J.; Yu, J.; Xu, Y.; Xing, Z.; Zhao, Y.; Wang, B.; Huang, X., Modular Access to Chiral Amines via Imine Reductase-Based Photoenzymatic Catalysis. J Am Chem Soc 2024, 146 (20), 14278–14286.

9. Sun, N.; Huang, J.; Qian, J.; Zhou, T. P.; Guo, J.; Tang, L.; Zhang, W.; Deng, Y.; Zhao, W.; Wu, G.; Liao, R. Z.; Chen, X.; Zhong, F.; Wu, Y., Enantioselective [2+2]-cycloadditions with triplet photoenzymes. Nature 2022, 611 (7937), 715–720.

10. Amer, M.; Wojcik, E. Z.; Sun, C.; Hoeven, R.; Hughes, J. M. X.; Faulkner, M.; Yunus, I. S.; Tait, S.; Johannissen, L. O.; Hardman, S. J. O.; Heyes, D. J.; Chen, G.-Q.; Smith, M. H.; Jones, P. R.; Toogood, H. S.; Scrutton, N. S., Low carbon strategies for sustainable bio-alkane gas production and renewable energy. Energy & Environmental Science 2020, 13 (6), 1818–1831.

11. Ma, Y.; Zhang, X.; Li, Y.; Li, P.; Hollmann, F.; Wang, Y., Production of fatty alcohols from non-edible oils by enzymatic cascade reactions. Sustainable Energy & Fuels 2020, 4 (8), 4232–4237.

12. Ma, Y.; Zhang, X.; Zhang, W.; Li, P.; Li, Y.; Hollmann, F.; Wang, Y., Photoenzymatic Production of Next Generation Biofuels from Natural Triglycerides Combining a Hydrolase and a Photodecarboxylase. ChemPhotoChem 2019, 4 (1), 39–44.

13. Cha, H. J.; Hwang, S. Y.; Lee, D. S.; Kumar, A. R.; Kwon, Y. U.; Voss, M.; Schuiten, E.; Bornscheuer, U. T.; Hollmann, F.; Oh, D. K.; Park, J. B., Whole-Cell Photoenzymatic Cascades to Synthesize Long-Chain Aliphatic Amines and Esters from Renewable Fatty Acids. Angew Chem Int Ed Engl 2020, 59 (18), 7024–7028.

14. Xu, J.; Hu, Y.; Fan, J.; Arkin, M.; Li, D.; Peng, Y.; Xu, W.; Lin, X.; Wu, Q., Light-Driven Kinetic Resolution of alpha-Functionalized Carboxylic Acids Enabled by an Engineered Fatty Acid Photodecarboxylase. Angew Chem Int Ed Engl 2019, 58 (25), 8474–8478.

15. Cheng, F.; Li, H.; Wu, D.-Y.; Li, J.-M.; Fan, Y.; Xue, Y.-P.; Zheng, Y.-G., Light-driven deracemization of phosphinothricin by engineered fatty acid photodecarboxylase on a gram scale. Green Chemistry 2020, 22 (20), 6815–6818.

16. Cha, H.-J.; Lee, H.-R.; Lee, J.; Oh, D.-K.; Park, J.-B., Chemoenzymatic Cascade Conversion of Linoleic Acid into a Secondary Fatty Alcohol Using a Combination of 13S-Lipoxygenase, Chemical Reduction, and a Photo-Activated Decarboxylase. ACS Sustainable Chemistry & Engineering 2021, 9 (32), 10837–10845.

17. Biegasiewicz, K. F.; Cooper, S. J.; Gao, X.; Oblinsky, D. G.; Kim, J. H.; Garfinkle, S. E.; Joyce, L. A.; Sandoval, B. A.; Scholes, G. D.; Hyster, T. K., Photoexcitation of flavoenzymes enables a stereoselective radical cyclization. Science 2019, 364 (6446), 1166–1169.

18. Fu, H.; Cao, J.; Qiao, T.; Qi, Y.; Charnock, S. J.; Garfinkle, S.; Hyster, T. K., An asymmetric sp(3)-sp(3) cross-electrophile coupling using ‘ene’-reductases. Nature 2022, 610 (7931), 302–307.

19. Fu, H.; Hyster, T. K., From Ground-State to Excited-State Activation Modes: Flavin-Dependent “Ene”-Reductases Catalyzed Non-natural Radical Reactions. Acc Chem Res 2024, 57 (9), 1446–1457.

20. Biegasiewicz, K. F.; Cooper, S. J.; Emmanuel, M. A.; Miller, D. C.; Hyster, T. K., Catalytic promiscuity enabled by photoredox catalysis in nicotinamide-dependent oxidoreductases. Nat Chem 2018, 10 (7), 770–775.

21. Black, M. J.; Biegasiewicz, K. F.; Meichan, A. J.; Oblinsky, D. G.; Kudisch, B.; Scholes, G. D.; Hyster, T. K., Asymmetric redox-neutral radical cyclization catalysed by flavin-dependent ‘ene’-reductases. Nat Chem 2020, 12 (1), 71–75.

22. Harrison, W.; Huang, X.; Zhao, H., Photobiocatalysis for Abiological Transformations. Acc Chem Res 2022, 55 (8), 1087–1096.

23. Huang, X.; Feng, J.; Cui, J.; Jiang, G.; Harrison, W.; Zang, X.; Zhou, J.; Wang, B.; Zhao, H., Photoinduced chemomimetic biocatalysis for enantioselective intermolecular radical conjugate addition. Nature Catalysis 2022, 5 (7), 586–593.

24. Ju, S.; Li, D.; Mai, B. K.; Liu, X.; Vallota-Eastman, A.; Wu, J.; Valentine, D. L.; Liu, P.; Yang, Y., Stereodivergent photobiocatalytic radical cyclization through the repurposing and directed evolution of fatty acid photodecarboxylases. Nat Chem 2024, 16 (8), 1339–1347.

25. Sorigue, D.; Hadjidemetriou, K.; Blangy, S.; Gotthard, G.; Bonvalet, A.; Coquelle, N.; Samire, P.; Aleksandrov, A.; Antonucci, L.; Benachir, A.; Boutet, S.; Byrdin, M.; Cammarata, M.; Carbajo, S.; Cuine, S.; Doak, R. B.; Foucar, L.; Gorel, A.; Grunbein, M.; Hartmann, E.; Hienerwadel, R.; Hilpert, M.; Kloos, M.; Lane, T. J.; Legeret, B.; Legrand, P.; Li-Beisson, Y.; Moulin, S. L. Y.; Nurizzo, D.; Peltier, G.; Schiro, G.; Shoeman, R. L.; Sliwa, M.; Solinas, X.; Zhuang, B.; Barends, T. R. M.; Colletier, J. P.; Joffre, M.; Royant, A.; Berthomieu, C.; Weik, M.; Domratcheva, T.; Brettel, K.; Vos, M. H.; Schlichting, I.; Arnoux, P.; Muller, P.; Beisson, F., Mechanism and dynamics of fatty acid photodecarboxylase. Science 2021, 372 (6538).

26. Heyes, D. J.; Lakavath, B.; Hardman, S. J. O.; Sakuma, M.; Hedison, T. M.; Scrutton, N. S., Photochemical Mechanism of Light-Driven Fatty Acid Photodecarboxylase. ACS Catal 2020, 10 (12), 6691–6696.

27. Spacey, H. J.; Healy, D.; Kalapothakis, J. M. D.; Ma, J.; Sakuma, M.; Barran, P. E.; Heyes, D. J.; Scrutton, N. S., Exploring the Increased Activity of the Blue Light-Dependent Photoenzyme Fatty Acid Photodecarboxylase under Violet Light. ACS Catal 2025, 15 (8), 6088–6097.

28. Lakavath, B.; Hedison, T. M.; Heyes, D. J.; Shanmugam, M.; Sakuma, M.; Hoeven, R.; Tilakaratna, V.; Scrutton, N. S., Radical-based photoinactivation of fatty acid photodecarboxylases. Anal Biochem 2020, 600, 113749.

29. Guo, X.; Xia, A.; Zhang, W.; Li, F.; Huang, Y.; Zhu, X.; Zhu, X.; Liao, Q., Anaerobic environment as an efficient approach to improve the photostability of fatty acid photodecarboxylase. Chinese Chemical Letters 2023, 34 (4).

30. Wright, P. J.; English, A. M., Scavenging with TEMPO* to identify peptide-and protein-based radicals by mass spectrometry: advantages of spin scavenging over spin trapping. J Am Chem Soc 2003, 125 (28), 8655–65.

31. Davies, M. J., The oxidative environment and protein damage. Biochim Biophys Acta 2005, 1703 (2), 93–109.

32. Wu, Y.; Paul, C. E.; Hollmann, F., Stabilisation of the Fatty Acid Decarboxylase from Chlorella variabilis by Caprylic Acid. Chembiochem 2021, 22 (14), 2420–2423.

33. Zhang, W.; Ma, M.; Huijbers, M. M. E.; Filonenko, G. A.; Pidko, E. A.; van Schie, M.; de Boer, S.; Burek, B. O.; Bloh, J. Z.; van Berkel, W. J. H.; Smith, W. A.; Hollmann, F., Hydrocarbon Synthesis via Photoenzymatic Decarboxylation of Carboxylic Acids. J Am Chem Soc 2019, 141 (7), 3116–3120.

34. Cardoso, D. R.; Franco, D. W.; Olsen, K.; Andersen, M. L.; Skibsted, L. H., Reactivity of bovine whey proteins, peptides, and amino acids toward triplet riboflavin as studied by laser flash photolysis. J Agric Food Chem 2004, 52 (21), 6602–6.

35. Aleksenko, V. A.; Anand, D.; Remeeva, A.; Nazarenko, V. V.; Gordeliy, V.; Jaeger, K.-E.; Krauss, U.; Gushchin, I., Phylogeny and Structure of Fatty Acid Photodecarboxylases and Glucose-Methanol-Choline Oxidoreductases. Catalysts 2020, 10 (9).

36. Meng, L.; Kalapothakis, J.; Ma, J.; Spacey, H.; Johannissen, L.; Beth Alen, A.; Heyes, D.; Scrutton, N.; Barran, P., Mechanism of photo-induced conformational changes in the photoenzyme fatty acid photodecarboxylase revealed by light footprinting ion mobility mass spectrometry. ChemRxiv 2025. https://chemrxiv.org/engage/chemrxiv/article-details/68e4e310bc2ac3a0e0b4273b

37. Shen, L.; Ji, H. F., A theoretical study on the quenching mechanisms of triplet state riboflavin by tryptophan and tyrosine. J Photochem Photobiol B 2008, 92 (1), 10–12.

38. Gagliardi, C. J.; Murphy, C. F.; Binstead, R. A.; Thorp, H. H.; Meyer, T. J., Concerted Electron–Proton Transfer (EPT) in the Oxidation of Cysteine. The Journal of Physical Chemistry C 2015, 119 (13), 7028–7038.

39. McCaslin, T. G.; Pagba, C. V.; Hwang, H.; Gumbart, J. C.; Chi, S. H.; Perry, J. W.; Barry, B. A., Tyrosine, cysteine, and proton coupled electron transfer in a ribonucleotide reductase-inspired beta hairpin maquette. Chem Commun (Camb) 2019, 55 (63), 9399–9402.

