## Supplementary material for "Triplet Quenching by Active Site Cysteine Residues Improves Photo-stability in Fatty Acid Photodecarboxylase"

**Experimental methods**

**Materials**

All materials were of analytical grade and were purchased from Sigma-Aldrich unless otherwise stated.

**Expression and purification of *Cv*FAP**

A truncated, codon-optimized *Cv*FAP sequence (residues 76-654) from *Chlorella variabilis* was synthesized (Geneart, Life Technologies) and cloned into the pET21a expression vector (C-terminal histidine-tagged thioredoxin). The plasmid was transformed into BL21 (DE3) *E. coli* cells for expression. Cells were grown in terrific broth (TB) medium at 37 °C and 200 rpm agitation to an OD_600_ of 1.0 before induction with 0.2 mM IPTG. Then the temperature was lowered to 18 °C and the cells were grown overnight at 180 rpm. Cells were harvested by centrifugation at 4 °C, 5000 rpm for 30 minutes.

All purification processes were performed at 4 °C and in a dark room. Cell pellets were resuspended in 50 mM Tris (pH 8.5), 300 mM NaCl, 5 % glycerol, 0.25 mg ml^-1^ lysozyme and 10 µg ml^-1^ DNase. Protease inhibitor cocktail tablets were added to the suspension to prevent proteolysis. Cells were lysed by sonication and clarified at 18000 rpm for 1 hour. The supernatant was loaded onto a HisTrap HP column pre-equilibrated with Buffer A (50 mM Tris (pH 8.5), 300 mM NaCl, and 5% glycerol). The column was washed with Buffer A and Buffer B (50 mM Tris (pH 8.5), 150 mM NaCl, 5% glycerol + 5 mM imidazole) to remove non-specifically bound proteins. Target protein was eluted with Buffer C (50 mM Tris (pH 8.5), 150 mM NaCl, 5% glycerol + 250 mM imidazole). Last-step purification was performed using size exclusion chromatography (HiLoad 200 16/60, GE Healthcare) with a buffer containing 50 mM Tris-HCl (pH 8.0), 150 mM NaCl, and 5% (v/v) glycerol. The purity of *Cv*FAP was confirmed by SDS-PAGE gels. The purified protein was frozen in liquid nitrogen and then stored at −80 °C before use. The concentration and quantity of flavin-bound protein were determined with extinction coefficients of 63,830 M⁻¹·cm⁻¹ at 280 nm for *Cv*FAP and 11,300 M⁻¹·cm⁻¹ at 450 nm for FAD cofactor released from the denatured protein with 0.2% SDS.

**Activity detection using flame-ionized detection gas chromatography**

*Cv*FAP was combined with substrates palmitic acid in Tris-HCl buffer pH 8.5 and the reaction was initiated using a 470 nm LED panel and quenched by placing samples in the dark at the desired time. Reactions were carried out in the well-plate at 350 rpm and 21 °C using the Eppendorf incubator. These samples were vigorously mixed with ethyl acetate with 0.1 % (v/v) *sec*-butyl benzene. Samples were centrifuged for 10 minutes using a benchtop centrifuge. Dried MgCl_2_ was added to the supernatant and centrifuged for a further 10 minutes. The supernatant was subsequently transferred and analyzed using gas chromatography with flame-ionized detection (GC-FID) with an HP-1 column (Agilent Technologies). The standard curve for the pentadecane was obtained to quantify the product in the reaction prior to detection of the samples (Figure S1).

**Steady state fluorescence measurements**

Data were acquired using Xe 900, Edinburgh Instrument Ltd (Livingston, UK) at room temperature by exciting the samples at 455 nm and recording the emission spectra from 460 to 650 nm. Samples contained ~10 µM *Cv*FAP in 50 mM Tris-HCl buffer pH 8.0, 150 mM NaCl and 5% glycerol. Excitation bandwidth was set to 0.3, the emission bandwidth was set to 2 and the iris was set at 20. Fluorescence spectra were recorded three times and measured with a dwell time of 1 s/nm. All measurements were performed in triplicate using a quartz cuvette with a 1 cm optical path length.

**Continuous wave electron paramagnetic resonance (CW-EPR) measurements**

EPR were performed with 150 μM *Cv*FAP. EPR samples were measured on a Bruker ELEXYSYS-E580 X-band EPR spectrometer. Microwave power was set to 33 dB (0.1 mW), and the modulation amplitude was set to 1 G. A time constant of 41 ms, a conversion time of 41 ms, and a sweep time of 84 s were used. The receiver gain was set to 60 dB with an average microwave frequency of 9.384 GHz. Samples were dissolved in 50 mM Tris-HCl (pH 8.0), 150 mM NaCl and 5% glycerol and all EPR measurements were performed in Suprail quartz-EPR tubes (Wilmad Labglass). All samples were frozen and stored in liquid nitrogen until measurements were performed at 20 K.

**Direct detection of hydrogen peroxide**

The assay was performed in a microtiter plate-based format. For each measurement, 100 μL sample containing 1 μM *Cv*FAP in 50 mM sodium phosphate buffer pH 8.0, 0.1 mM 10-acetyl-3,7-dihydroxyphenoxazine (ADHP), and 0.2 U mL^−1^ horseradish peroxidase (HRP) was illuminated under blue light and detected after the illumination. For comparison, solutions with defined hydrogen peroxide concentrations were measured with the same assay to obtain the standard curve. Hydrogen peroxide production was followed by absorbance measurement at 560 nm (FLUOstar platereader).

**Laser photoexcitation measurements**

Data were acquired using an image-intensified CCD camera (Andor Technologies) of an LP980 laser flash photolysis instrument (Edinburgh Instruments Ltd) upon excitation with a laser pulse (6-8 ns) from a Q-switched Nd-YAG laser (NT432, EKSPLA) in a cuvette of 1 cm pathlength. Detection of the red-shifted flavin intermediate and triplet excited state was performed using the same nanosecond laser flash photolysis system. Kinetic absorption transients were recorded at 515 nm or 690 nm with the detection system (comprising probe light, sample, monochromator and photomultiplier) at right angles to the incident laser beam. Samples contained 50 µM *Cv*FAP with 300 µM palmitic acid in 50 mM Tris-HCl buffer containing 10 % DMSO at pH 8.5. The reaction was driven using laser pulses 455nm (~17 mJ) to maximize the signal intensity. Time constants were observed from the average of at least five time-dependent absorption measurements by fitting them to an exponential function using the L900 software (Edinburgh Instruments Ltd).

**Kinetic modelling of FAP photocatalysis and photoinactivation**

Differential equations of the kinetic models employed in this study were solved analytically and presented below here. Non-linear curve fitting was performed by Origin(Pro) 2022b (OriginLab Corporation, Northampton, MA, USA). The fitting functions were expressed in LabTalk; non-linear curve fitting was performed iteratively using the Levenberg-Marquardt algorithm and terminated when the reduction in $\chi^{2}$ dropped below 10^-9^. The inactivation or activation rate coefficients, $k_{i}$ and $\beta$, were obtained directly from fitted parameters, and the catalytic rate coefficients $\alpha_{S}k$ could be obtained by simple algebraic manipulations of fitted parameters.

Here is our kinetics modelling. Given that catalysis and photo-inactivation are irreversible steps in the mechanism, assuming that either the protein:product complex or the dissociated substrate-free and product forms are favored, the substrate is converted to product irreversibly when it encounters the protein, with $C$ being the concentration of active catalyst (protein) and $S$ is the concentration of substrate:

$$\frac{dS}{dt}=-kv_{I}Cf\left( S \right)$$

[Equation 1a]

where $k$ is the reaction rate constant per concentration unit per photon current unit per second and $v_{I}$is the light intensity, $I$, dependence of the reaction, with the same photon current units. $f\left( S \right)$ describes the dependence of the reaction rate on substrate concentration, and we have empirically verified that a logistic function adequately describes this relationship, i.e. $f\left( S \right)=a_{s}\frac{e^{\kappa S}-1}{e^{\kappa S}+1}$ where $a_{s}$ is the activity of the substrate, a constant that can be absorbed into the reaction rate coefficient, and $\kappa$ determines the shape of the substrate scaling term; with units of 1/[concentration], it can be understood as a binding equilibrium constant, but one that is valid in dynamic reaction conditions. $a_{s}$ and $\kappa$ have been determined by fitting the dependence of the experimentally observed reaction rates to the substrate concentration (Figure S1).

Equation 1a leads to an exact expression for the substrate as a function of time:

$$S\left( t \right)=\frac{2}{\kappa}\sinh^{-1}\left[ \left( \sinh\left( \frac{\kappa}{2}S_{0} \right) \right)e^{-\kappa a_{s}k\int_{0}^{t} C(t^{'})dt^{'}} \right]$$

[Equation 1b]

Therefore, an exact expression is obtained with a suitable choice of $C\left( t \right)$.

The inactivation profile of those variants suggests a mechanism whereby two protein states photo-inactivate with rate coefficients $k_{i_{1}}v_{I}$ and $k_{i_{2}}v_{I}$ respectively, $v_{I}$ being the dependence of the inactivation rate coefficient on light intensity and $k_{i}$ the effective cross section of the photo-inactivation reaction:

$$\frac{dC_{1}}{dt}=-k_{i_{1}}v_{I}C_{1}$$

$$\frac{dC_{2}}{dt}=-k_{i_{2}}v_{I}C_{2}$$

$$C=C_{1}+C_{2}$$

[Equation 2a-c]

Under the assumption that the enzyme states $C_{1}$ and $C_{2}$ are not in rapid equilibrium. These combine to a solution:

$$C\left( t \right)=C_{1_{0}}e^{-k_{i_{1}}v_{I}t}+C_{2_{0}}e^{-k_{i_{2}}v_{I}t}$$

[Equation 2d]

Surprisingly, an early photo-activation step is observed in those variants. The dissociation of fatty acids from FAP is inefficient; thus, this early step may correspond to the turnover of natively bound substrate, or the migration of the fatty acid from the allosteric binding site to the active site, or the photo-induced dissociation of transient oligomers. The process will be of the form:

$$C_{1}\underset{\to}{\beta}C$$

$$\frac{dC_{1}}{dt}=-\beta C_{1}$$

$$\frac{dC}{dt}=\beta C_{1}-k_{i}v_{I}C$$

$$C\left( t \right)=\left( C_{0}+C_{1_{0}}\gamma\right)e^{-k_{i}v_{I}t}-C_{1_{0}}\gamma e^{-\beta t}$$

$$\gamma=\frac{1}{1-\frac{k_{i}v_{I}}{\beta}}$$

[Equation 3a-e]

Here, $C_{1}$ is the precursor state of the active enzyme $C$, which transitions to the active state with rate coefficient $\beta$. Substituting Equation 3d into Equation 1b and integrating we obtain:

$$S\left( t \right)=\frac{2}{\kappa}\sinh^{-1}\left[ \left( \sinh\left( \frac{\kappa}{2}S_{0} \right) \right)\text{exp}\left( -\kappa a_{s}\frac{k}{k_{i}}C_{0}\tau\right) \right]$$

$$\tau=\left( \left( 1+\gamma\frac{{C_{1}}_{0}}{C_{0}} \right)\left( 1-e^{{-k}_{i}v_{I}t} \right)-\frac{\gamma k_{i}v_{I}{C_{1}}_{0}}{\beta C_{0}}\left( 1-e^{-\beta t} \right) \right)$$

$$\gamma=\frac{1}{1-\frac{k_{i}v_{I}}{\beta}}$$

[Equation 4]

Equation 4 describes the density of substrate as a function of time. Note that, unlike catalysis by a stable catalyst, at long times a part of the substrate does not react despite the fact that the photodecarboxylation is treated as irreversible. The equilibrium substrate density is:

$$S_{eq}=\frac{2}{\kappa}\sinh^{-1}\left[ \left( \sinh\left( \frac{\kappa}{2}S_{0} \right) \right)\text{exp}\left( -\kappa a_{s}\frac{k}{k_{i}}C_{0} \right) \right]$$

[Equation 5]

Therefore, the reaction yield $S_{0}-S_{eq}$ is related to the dimensionless quantity $a_{s}\kappa\frac{k}{k_{i}}C_{0}$, which characterizes the kinetics of catalysis competing with photo-inactivation.

**Molecular Dynamics Simulations**

Molecular Dynamics simulations were performed using the drMD software package^1^, which uses OpenMM^2^ to perform MD simulations. Prior to using drMD, protons were added to the FAD cofactor using Pymol^3^. AMBER compatible forcefield parameters were created for the FAD cofactor using a combination of antechamber and parmchk, from the AmberTools suite^4^. All proteins were protonated using software pdb2pqr^5-6^ which uses ProPKA to calculate per-residue proton affinities^7-8^. Protein residues were protonated at a pH of 7.4. This process also automatically creates disulfide bonds as appropriate. All proteins were placed in a cubic solvation box with a 10 Å buffer between the protein and the nearest edge of the box. The system was treated using periodic boundary conditions. Approximately 30000 TIP3P water molecules were added to the solvation box. Sodium ions were added to the box to balance the charge of the system. All protein residues were parameterised using the AMBER ff19SB and forcefield^9^. These parameters were prepared using tleap from the Ambertools suite^4^. Simulations were performed in explicit solvent. All water molecules parameterised using the TIP3P model^10^. Any ions in our system were treated using parameters calculated to complement the TIP3P water model^11^.

For all systems, the following simulation protocol was used in triplicate: Initially, an energy minimisation step was performed using the steepest descent method. This energy minimisation step was performed for 1000 steps, or until it reached convergence. Next, a simulation was performed using the canonical (NVT) ensemble. This simulation was performed for 100 picoseconds. This was followed by a 100-picosecond simulation performed under the isothermal-isobaric (NpT) ensemble. For the two previously described simulation steps, restraints were applied to the positions of all protein and ligand atoms with a force constant of 1000 kJ/mol/nm^2.^ Next a short (10 picoseconds) simulation was performed under the NpT ensemble using a timestep of 0.5 femtoseconds, the purpose of this step was to allow the system to relax after the removal of the position restraints. A longer (5 nanoseconds) equilibration was then performed under the NpT ensemble. Production MD simulations were performed for 50 nanoseconds under the NpT ensemble. All simulation steps were performed at 300 K.

With the exception of the step run with a timestep of 0.5 femtoseconds, the mass of hydrogen atoms was set to 4.03036 amu, this allowed these simulation steps to be performed using a timestep of 4 femtoseconds, resulting in an increase in simulation speed. All simulations were performed using the Langevin Middle Integrator^12^ this was used to enforce a constant temperature in each simulation. For simulations run under the NpT ensemble, the Monte-Carlo barostat was used to enforce a constant pressure of 1 atm. In all simulations, long-range Coulombic interactions were modelled using the Particle-Mesh Ewald (PME) method, with a 10 Å cutoff distance. In all simulations, constraints were applied to bonds between hydrogen atoms and heavy atoms.

Analysis of molecular dynamics trajectories was performed in the MDAnalysis python package^13-14^.

**QM/MM calculations**

Starting structures for QM/MM calculations were generated from the last structure of 100 ns MD simulations performed in Gromacs 2020.3^15-16^. The system was set up in the same manner as described in the MD section, after which the Amber toplogy and structure files were converted into Gromacs format using ParmEd^17^. MD simulations were then performed using LINCS bond constraints^18^ applied to all bonds involving hydrogen, Verlet integration, 10 Å cut-offs for Coulombic and van der Waals interactions, Particle Mesh Ewald for long-range electrostatics, the velocity-rescaling modified Berendsen thermostat^19^ (300 K), the Parrinello-Rahman barostat^20^ (1 bar), periodic boundary and a 2 fs timestep. Simulations were run according to the following protocol: (i) energy minimisation (ii) 1 ns constant volume (NVT) equilibration of the solvent with 10 kJ mol^-1^ Å^-2^ force constant position restraints on the protein non-hydrogen atoms; (iii) 1 ns constant pressure (NPT) equilibration of the solvent with the same position restraints; (iv) 1 ns constant pressure (NPT) equilibration of the solvent with a 1 kJ mol^-1^ Å^-2^ force constant; (v) 1 ns unrestrained MD simulations. The structure for the QM/MM calculations was defined as the protein, FAD and all water molecules with at least one atom within 30 Å of the FAD N5 (Figure SX1), for a total of 13,901 atoms for A158C, and 13820 atoms for A171C.

QM/MM calculations were performed in Gaussian16 rev C.01^21^ with the QM-MM electrostatic interaction treated by electronic embedding. The QM region was treated at the ωB97-XD/6-31G(d,p) level of theory and was defined as the FAD and the sidechain of either C158 (for A158C), C622 (for G622C) or C171 (for A171C), and the sidechain of D139 for A158C. For A171C the nearest water molecule bridging the C171 thiol and FAD N5 was also included so that H-transfer *via* this water molecule could also be modelled. In each case a link atom was positioned between the Cα(MM) and Cβ(QM) of the cysteine and D139 for A158C. The majority of the system was kept fixed during the calculations except for a free region defined as follows: for A158C and G622C, all residues or water molecules with at least one atom within 10 Å of N5 of FAD or S of C158/C622 (1142 and 1160 atoms, respectively); for A171C all residues or water molecules with at least one atom within 12 Å of FAD N5 (1181 atoms). Each model was first energy minimised in the ground state (s0), and this geometry was then energy minimised as a triplet with a spin multiplicity of 3 on the QM region. A relaxed scan was then performed along a reaction coordinate *z* for proton transfer. For direct H-transfer between thiol and the proton acceptor (N5 for A171C, N1 for G622C and sidechain carboxylate oxygen for D139C), this was defined as the difference between the lengths of the breaking and forming bonds. For H-transfer *via* bridging water molecule in A171C, the reaction coordinate was defined as the average difference between the two breaking and forming bonds. For G622C the potential energy scan suggests an adiabatic PCET (hydrogen atom transfer, HAT), *ie.* the electron transfer is concurrent with the proton transfer. However, for A171C and D139C scans are consistent with nonadiabatic PCET and the system jumps from the reactant state [^3^FAD + SH] electronic surface to the product state ^3^[FADH· + S·] electronic surface. After energy minimizing the PCET product, potential energy scans were then carried out in the reverse direction to obtain the energetic crossing point for the H-coordinate along each electronic surface (*cf.* the transition state in Marcus transition theory).

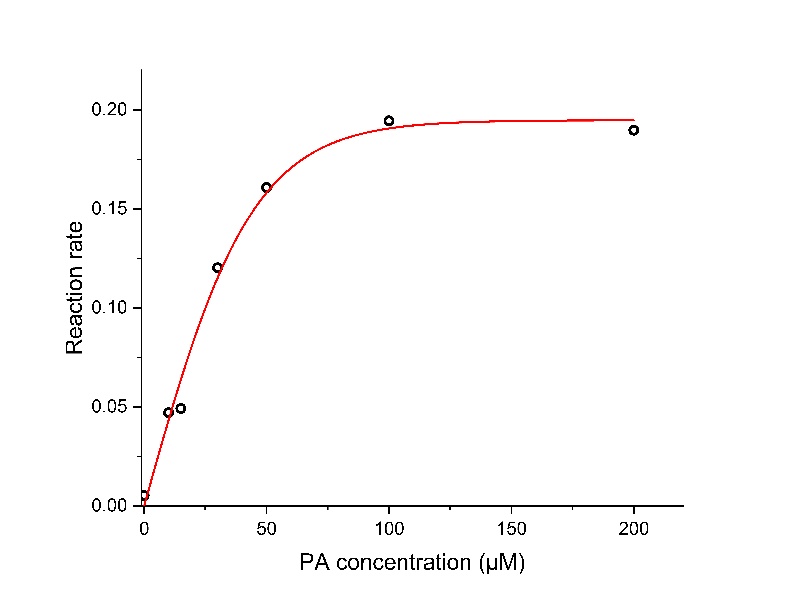

**Figure S1.** Initial reaction rate of FAP with various concentration of palmitic acid determined by the stopped-flow based coupled assay under blue light.

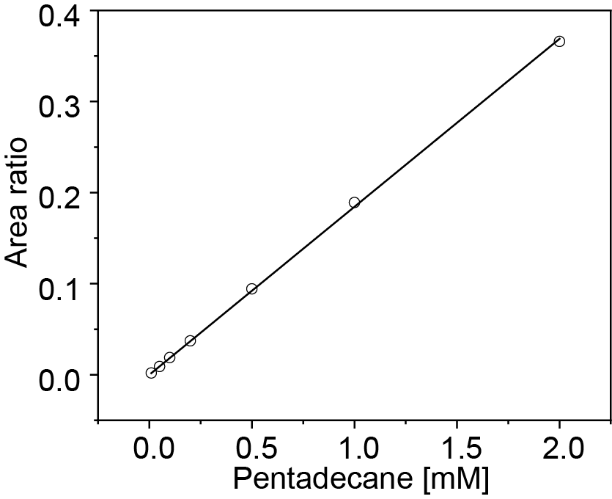

**Figure S2.** Standard curve for pentadecane detected by GC-FID using HP-1 column. The area ratio in the figure is the peak area ratio of pentadecane to sec-butyl benzene.

**Figure**
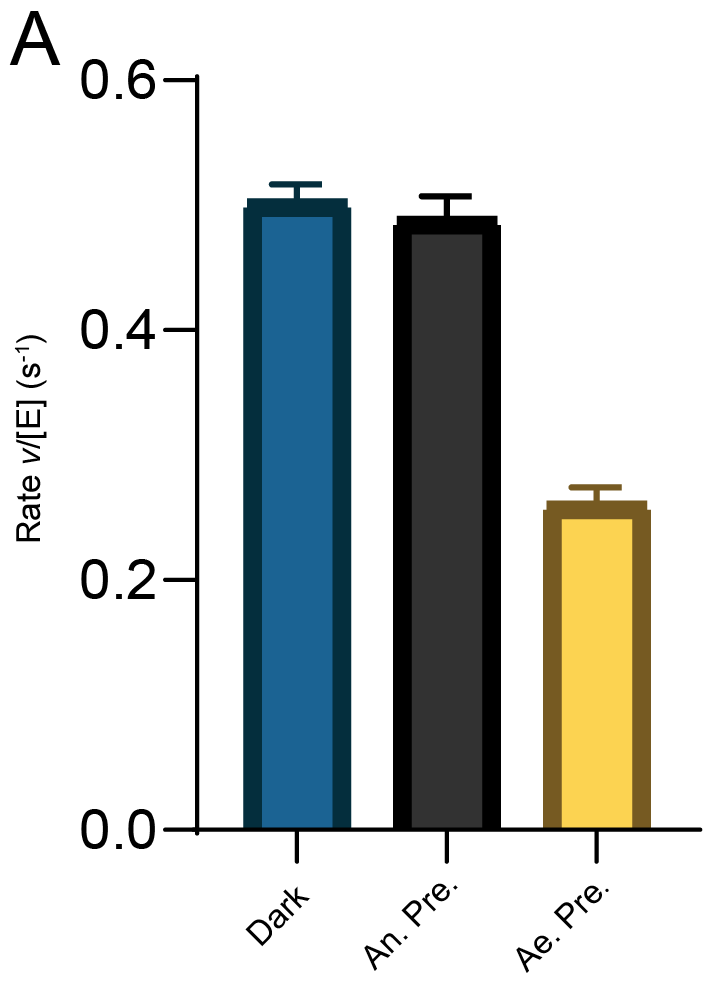
**S3.** Activity of wild-type FAP before and after pre-illumination of the substrate-free enzyme with 1000 μmol m^-2^s^-1^ blue light under aerobic and anaerobic conditions at room temperature. The initial rate of activity was measured using the stopped-flow based coupled assay to measure the rate of CO_2_ formation.

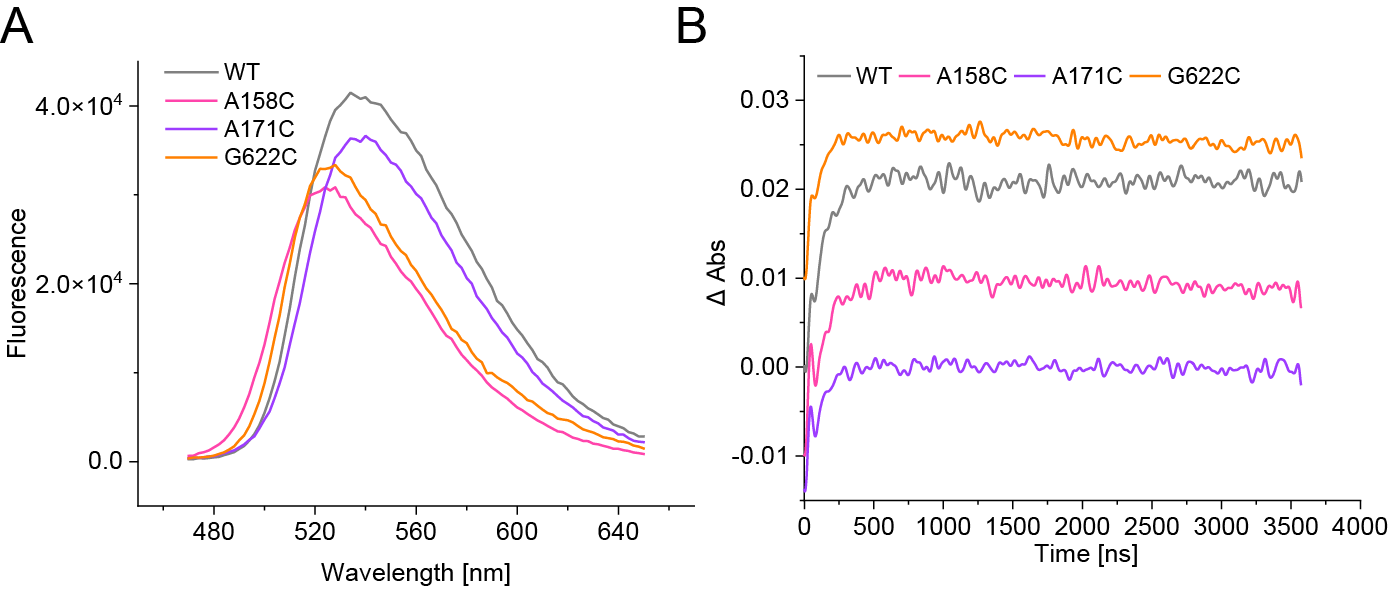
**Figure S4.** Laser photoexcitation experiments probing formation of red-shifted species at 515 nm in wild-type and double Cys variants of FAP. Transient absorption changes were measured over 4 μs for samples containing 40μM FAP and 500μM palmitic acid and 10% DMSO after excitation with a laser pulse at 450 nm (~15 mJ).

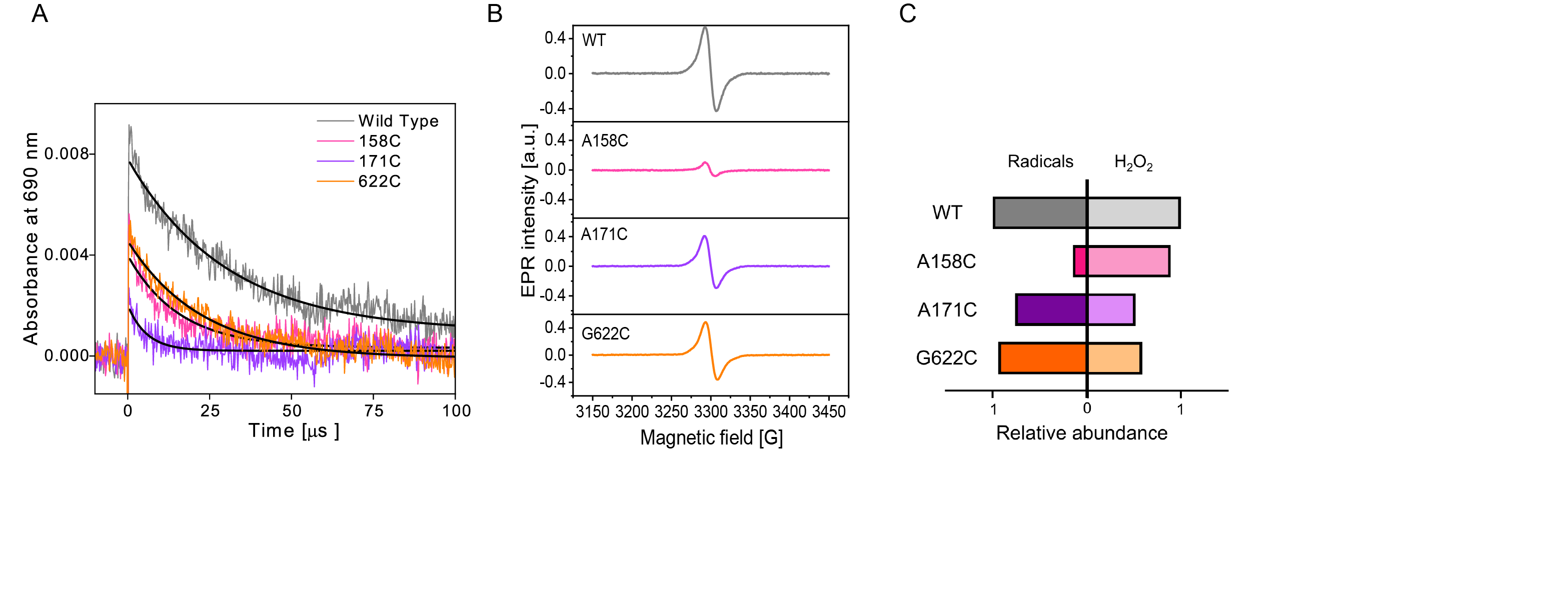

**Figure S5.** Detection of radical accumulation by EPR spectroscopy. Continuous-wave electron paramagnetic resonance spectroscopy (CW-EPR) showing the formation of radical species when 150 μM samples were illuminated for only 15 s with a 455 nm LED of 1000 μmol m^-2^s^-1^.

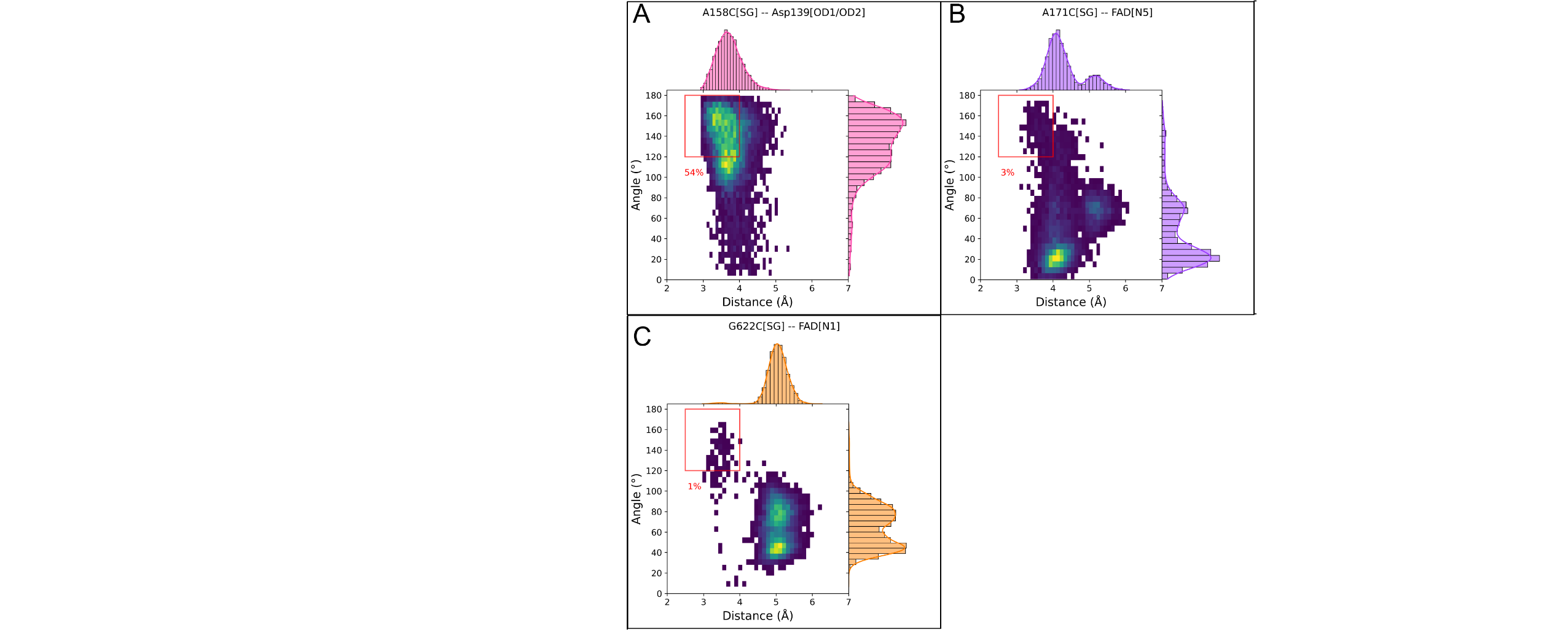

**Figure S6.** Plots distributions of hydrogen bond distances and angles throughout MD simulations for: The interaction of A158C with Asp139 (A), the interaction of A171C with FAD N_5_ (B) and the interaction of A171C with Tyr466 (C). Histograms of inter-heteroatom distances are shown along the x-axis. Histograms of the hydrogen bond angle (defined by Cys-SG–Cys-HG- Heteroatom) are shown along the y-axis. On each plot, a red box shows the region of distance-angle space in which a hydrogen bond is formed, above this the percentage of structures in MD simulations where hydrogen bonds are formed is annotated.

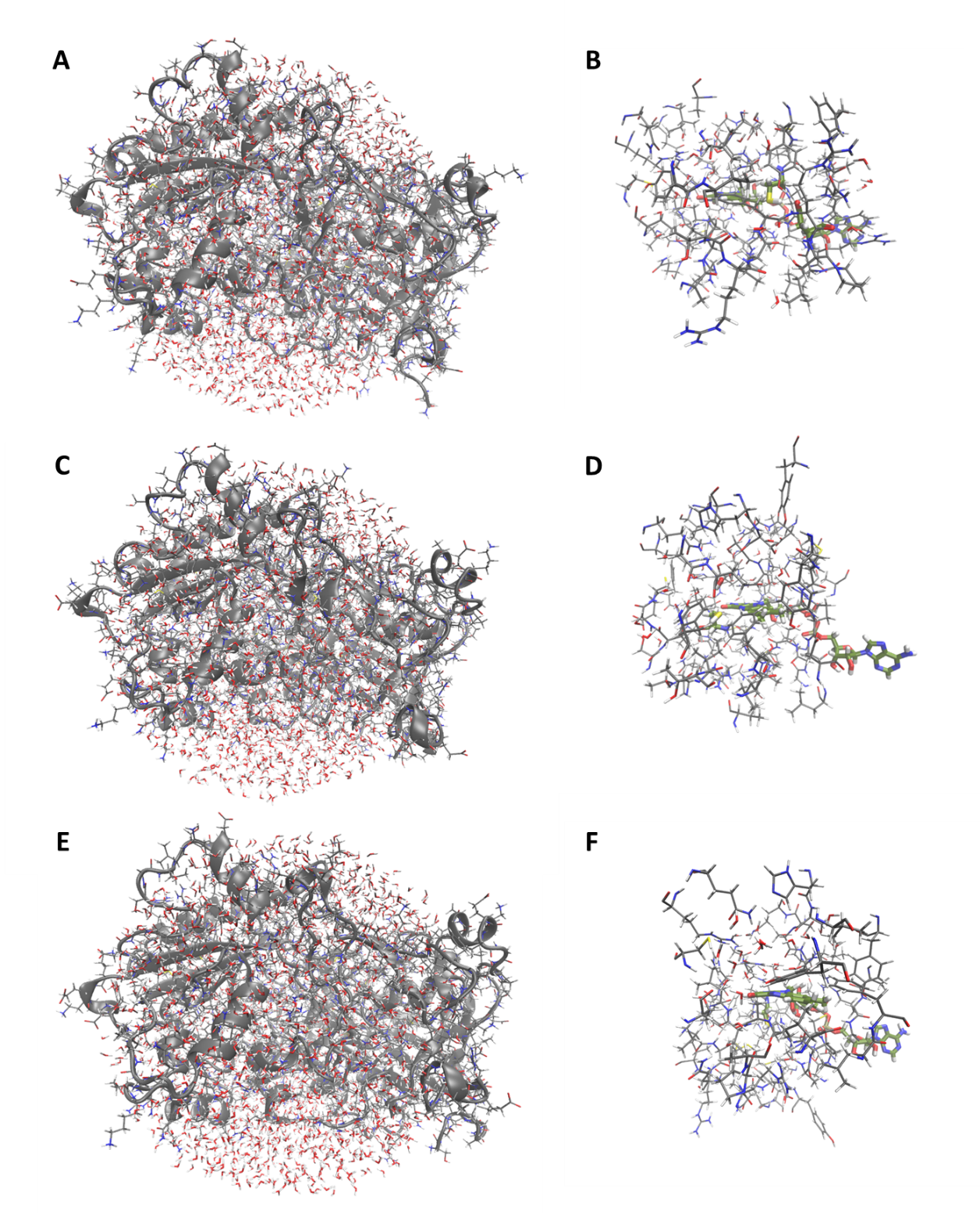

**Figure S7.** QM/MM models of the three cvFAP variants A158C (A,B), A171C (C,D) and G622C (E,F). The full models (A,C,E) are shown with the protein as a ribbon and the protein, FAD and water as sticks; the regions of the models that are not fixed during the calculations (B,D,F) are shown with the MM region as thin lines and the QM region as thicker sticks.

**Table S1**: Variants of CvFAP containing a single arginine or cysteine residue in C432S or R451A variants. All of these double variants are inactive.

| Mutation site | Symbol |
| --- | --- |
| Wild type | WT |
| I130 | I130C C432S |
| F134 | F134C C432S |
| T169 | T169C C432S |
| A171 | A171C C432S |
| T430 | T430C C432S |
| V463 | V463C C432S |
| Y466 | Y466C C432S |
| Q486 | Q486C C432S |
| H572 | H572C C432S |
| S573 | S573C C432S |
| S574 | S574C C432S |
| N575 | N575C C432S |
| I130 | I130R R451A |
| D139 | D139R R451A |
| A158 | A158R R451A |
| L173 | L173R R451A |
| V463 | V463R R451A |
| I488 | I488R R451A |

**Table S2**: Variants of CvFAP containing a single arginine or cysteine residue in wild-type FAP. All of these variants are inactive.

| Mutation site | Symbol |
| --- | --- |
| F134 | F134C |
| T430 | T430C |
| Q486 | Q486C |
| N575 | N575C |
| Q620 | Q620C |
| D139 | D139R |
| A158 | A158R |
| L173 | L173R |
| V463 | V463R |
| I488 | I488R |
| N575 | N575R |

**Table S3: Summary of QM/MM calculations of PCET between cysteine and FAD: energies and spin densities and partial charges for the ground, excited triplet and semiquinone FAD states.**

| Chemical species | *ΔE* / kJ mol^-1^ | *q*(iso)^1^ | *σ*(iso)^2^ |
| --- | --- | --- | --- |
| A158C | | | |
| ^1^FAD + ^1^cys-H | 0 | -0.139 |  |
| ^3^FAD* + ^1^cys-H | 191 | -0.145 | 1.97 |
| ^3^[FAD· + cys·] | 169 | -1.07 | 0.993 |
| A171C | | | |
| ^1^FAD + ^1^cys-H (min) | -6.62 | -0.0852 |  |
| ^1^FAD + ^1^cys-H | 0 | -0.0764 |  |
| ^3^FAD* + ^1^cys-H | 195 | -0.0623 | 1.96 |
| ^3^[FAD-H· + cys·] | 157 | -0.436 | 0.990 |

^1^ Partial charge on FAD isoalloxazine (sum of Mulliken atomic charges)

^2^ Spin multiplicity on FAD isoalloxazine (sum of Mulliken multiplicities)

REFERENCES

1.	Shrimpton-Phoenix, E.; Notari, E.; Kluonis, T.; Wood, C. W., drMD: Molecular Dynamics for Experimentalists. *J Mol Biol* **2025,** *437* (15), 168918.

2.	Eastman, P.; Swails, J.; Chodera, J. D.; McGibbon, R. T.; Zhao, Y.; Beauchamp, K. A.; Wang, L. P.; Simmonett, A. C.; Harrigan, M. P.; Stern, C. D.; Wiewiora, R. P.; Brooks, B. R.; Pande, V. S., OpenMM 7: Rapid development of high performance algorithms for molecular dynamics. *PLoS Comput Biol* **2017,** *13* (7), e1005659.

3.	Schrödinger, L. *The PyMOL Molecular Graphics System, Version 1.8*, Schrödinger, LLC: New York, 2015.

4.	Case, D. A.; Aktulga, H. M.; Belfon, K.; Cerutti, D. S.; Cisneros, G. A.; Cruzeiro, V. W. D.; Forouzesh, N.; Giese, T. J.; Götz, A. W.; Gohlke, H.; Izadi, S.; Kasavajhala, K.; Kaymak, M. C.; King, E.; Kurtzman, T.; Lee, T.-S.; Li, P.; Liu, J.; Luchko, T.; Luo, R.; Manathunga, M.; Machado, M. R.; Nguyen, H. M.; O’Hearn, K. A.; Onufriev, A. V.; Pan, F.; Pantano, S.; Qi, R.; Rahnamoun, A.; Risheh, A.; Schott-Verdugo, S.; Shajan, A.; Swails, J.; Wang, J.; Wei, H.; Wu, X.; Wu, Y.; Zhang, S.; Zhao, S.; Zhu, Q.; Cheatham, T. E., III; Roe, D. R.; Roitberg, A.; Simmerling, C.; York, D. M.; Nagan, M. C.; Merz, K. M., Jr., AmberTools. *Journal of Chemical Information and Modeling* **2023,** *63* (20), 6183-6191.

5.	Dolinsky, T. J.; Czodrowski, P.; Li, H.; Nielsen, J. E.; Jensen, J. H.; Klebe, G.; Baker, N. A., PDB2PQR: expanding and upgrading automated preparation of biomolecular structures for molecular simulations. *Nucleic Acids Res* **2007,** *35* (Web Server issue), W522-5.

6.	Dolinsky, T. J.; Nielsen, J. E.; McCammon, J. A.; Baker, N. A., PDB2PQR: an automated pipeline for the setup of Poisson-Boltzmann electrostatics calculations. *Nucleic Acids Res* **2004,** *32* (Web Server issue), W665-7.

7.	Olsson, M. H.; Sondergaard, C. R.; Rostkowski, M.; Jensen, J. H., PROPKA3: Consistent Treatment of Internal and Surface Residues in Empirical pKa Predictions. *J Chem Theory Comput* **2011,** *7* (2), 525-37.

8.	Sondergaard, C. R.; Olsson, M. H.; Rostkowski, M.; Jensen, J. H., Improved Treatment of Ligands and Coupling Effects in Empirical Calculation and Rationalization of pKa Values. *J Chem Theory Comput* **2011,** *7* (7), 2284-95.

9.	Tian, C.; Kasavajhala, K.; Belfon, K. A. A.; Raguette, L.; Huang, H.; Migues, A. N.; Bickel, J.; Wang, Y.; Pincay, J.; Wu, Q.; Simmerling, C., ff19SB: Amino-Acid-Specific Protein Backbone Parameters Trained against Quantum Mechanics Energy Surfaces in Solution. *J Chem Theory Comput* **2020,** *16* (1), 528-552.

10.	Neria, E.; Fischer, S.; Karplus, M., Simulation of activation free energies in molecular systems. *The Journal of Chemical Physics* **1996,** *105* (5), 1902-1921.

11.	Li, P.; Song, L. F.; Merz, K. M., Jr., Systematic Parameterization of Monovalent Ions Employing the Nonbonded Model. *J Chem Theory Comput* **2015,** *11* (4), 1645-57.

12.	Zhang, Z.; Liu, X.; Yan, K.; Tuckerman, M. E.; Liu, J., Unified Efficient Thermostat Scheme for the Canonical Ensemble with Holonomic or Isokinetic Constraints via Molecular Dynamics. *J Phys Chem A* **2019,** *123* (28), 6056-6079.

13.	Michaud-Agrawal, N.; Denning, E. J.; Woolf, T. B.; Beckstein, O., MDAnalysis: a toolkit for the analysis of molecular dynamics simulations. *J Comput Chem* **2011,** *32* (10), 2319-27.

14.	Gowers, R. J.; Linke, M.; Barnoud, J.; Reddy, T. J. E.; Melo, M. N.; Seyler, S. L.; Domanski, J.; Dotson, D. L.; Buchouxk, S.; Kenney, I. M.; Beckstein, O., MDAnalysis: A Python Package for the Rapid Analysis of Molecular Dynamics Simulations. In *Proceedings of the Python in Science Conference (SciPy)*, 2016.

15.	Páll, S.; Abraham, M. J.; Kutzner, C.; Hess, B.; Lindahl, E. In *Tackling Exascale Software Challenges in Molecular Dynamics Simulations with GROMACS*, Solving Software Challenges for Exascale, Cham, 2015//; Markidis, S.; Laure, E., Eds. Springer International Publishing: Cham, 2015; pp 3-27.

16.	Abraham, M. J.; Murtola, T.; Schulz, R.; Páll, S.; Smith, J. C.; Hess, B.; Lindahl, E., GROMACS: High performance molecular simulations through multi-level parallelism from laptops to supercomputers. *SoftwareX* **2015,** *1-2*, 19-25.

17.	Swails, J.; Hernandez, C.; Mobley, D. L.; Nguyen, H.; Wang, L.-P.; Janowski, P. ParmEd. <https://github.com/ParmEd/ParmEd> (accessed October 2, 2025).

18.	Hess, B.; Bekker, H.; Berendsen, H. J. C.; Fraaije, J. G. E. M., LINCS: A linear constraint solver for molecular simulations. *Journal of Computational Chemistry* **1997,** *18* (12), 1463-1472.

19.	Bussi, G.; Donadio, D.; Parrinello, M., Canonical sampling through velocity rescaling. *The Journal of Chemical Physics* **2007,** *126* (1), 014101.

20.	Aoki, K. M.; Yonezawa, F., Constant-pressure molecular-dynamics simulations of the crystal-smectic transition in systems of soft parallel spherocylinders. *Phys Rev A* **1992,** *46* (10), 6541-6549.

21.	Frisch, M. J.; Trucks, G. W.; Schlegel, H. B.; Scuseria, G. E.; Robb, M. A.; Cheeseman, J. R.; Scalmani, G.; Barone, V.; Petersson, G. A.; Nakatsuji, H.; Li, X.; Caricato, M.; Marenich, A. V.; Bloino, J.; Janesko, B. G.; Gomperts, R.; Mennucci, B.; Hratchian, H. P.; Ortiz, J. V.; Izmaylov, A. F.; Sonnenberg, J. L.; Williams; Ding, F.; Lipparini, F.; Egidi, F.; Goings, J.; Peng, B.; Petrone, A.; Henderson, T.; Ranasinghe, D.; Zakrzewski, V. G.; Gao, J.; Rega, N.; Zheng, G.; Liang, W.; Hada, M.; Ehara, M.; Toyota, K.; Fukuda, R.; Hasegawa, J.; Ishida, M.; Nakajima, T.; Honda, Y.; Kitao, O.; Nakai, H.; Vreven, T.; Throssell, K.; Montgomery Jr., J. A.; Peralta, J. E.; Ogliaro, F.; Bearpark, M. J.; Heyd, J. J.; Brothers, E. N.; Kudin, K. N.; Staroverov, V. N.; Keith, T. A.; Kobayashi, R.; Normand, J.; Raghavachari, K.; Rendell, A. P.; Burant, J. C.; Iyengar, S. S.; Tomasi, J.; Cossi, M.; Millam, J. M.; Klene, M.; Adamo, C.; Cammi, R.; Ochterski, J. W.; Martin, R. L.; Morokuma, K.; Farkas, O.; Foresman, J. B.; Fox, D. J. *Gaussian 16 Rev. C.01*, Wallingford, CT, 2016.
